# Self-determination motivation theory in R: The software package S D T

**DOI:** 10.1101/510115

**Authors:** Ali Ünlü

## Abstract

This paper presents the technical details of the software package `SDT` in the `R` computing and graphics environment, implementing a convex quadratic program that was recently proposed in the literature on self-determination theory of human motivation. Three main features are addressed, with their accompanying code for computation in R: first, the application of the quadratic program and corresponding code for the analysis of the extent of motivation internalization or externalization; second, for exploring the simplex structure assumption of motivation; and third, for adjusting the confounded scoring protocol, called the self-determination or relative autonomy index, to account for the mixture of internal motivation and external motivation. We describe the functions of the `R` package `SDT`. The computations are demonstrated with example data accompanying the package, so researchers can run the methodology on their own datasets.

## 1. INTRODUCTION

This paper discusses the software package `SDT` for self-determination theory (SDT) measures in the `R` computing and graphics environment (The `R` Core Team, 2016). The package is available at no cost from the Comprehensive R Archive Network (CRAN), http://CRAN.R-project.org/package=SDT. SDT was introduced by Deci and Ryan (1985, 2000, 2002) and provides a theoretical framework for studying motivation. With SDT, researchers can describe the motivational basis of human behavior. On the one hand, there are the extrinsic forces acting on people (e.g., grades or evaluations), and on the other, there are the intrinsic motives inherent in humans (e.g., interests or curiosity). The general aim of SDT is to study the interplay between these extrinsic and intrinsic factors. In particular, types of motivation were postulated in SDT (Figure 1). For the introjected regulation and the identified regulation types of extrinsic motivation, their internalizations were delineated as *“somewhat external”* and *“somewhat internal”*, respectively. These notions remained undetermined in `SDT` (Ünlü & Dettweiler, 2015; Ünlü, 2016). The basic concepts of `SDT` are explained in Section 2.

**Figure 1.**
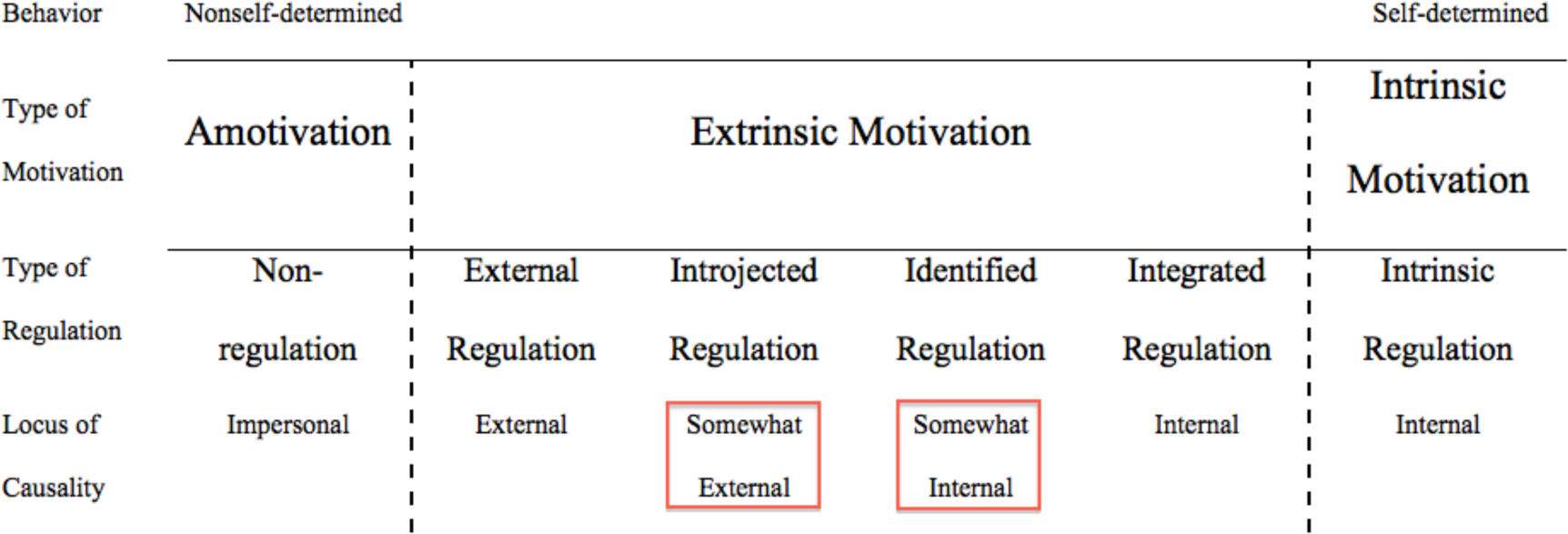
“The self-determination continuum, showing the motivational, self-regulatory, and perceived locus of causality bases of behaviors that vary in the degree to which they are self-determined” (Deci & Ryan, 2000, p. 237).

What makes this software package so exciting is threefold. First, the function `internalization` of the package `SDT` implements the constrained regression analysis approach proposed by Ünlü and Dettweiler (2015). Based on this approach, the vaguely expressed intermediate motivations can be estimated from data. The approach can be extended to simplex structure analysis, that is, for validating whether or not motivation regulations theoretically closer to one another are more strongly interrelated. Second, simplex structure analysis in `R` is realized with the function `simplex` of the package `SDT`. Third, the function `sdi` of the package provides the popular self-determination or relative autonomy index (SDI or RAI), in the common and adjusted variants. This is a scoring protocol that aggregates the subscale scores to imply an overall informative measure. The original SDI or RAI index is confounded. Generally, it does not accommodate biasing effects on the index value that may result from mixed internal and external motivation. Because of this, the function `sdi` also implements an adjusted scoring protocol variant of this measure, as discussed by Ünlü (2016). Thus, this package gives the user the ability to calculate adjusted SDI scores for all participants. Examples of this can be found in Section 5. In addition, the package `SDT` provides `plot, print`, and summary methods for objects of specific classes for conveniently graphing, printing, and outlining the results obtained from `SDT` analyses.

The paper has the following structure. In Section 2, the theory of self-determination is reviewed. In Section 3, we recapitulate the issues of motivation internalization, motivation simplex structure, and of adjusting the SDI or RAI measure for mixed internal and external motivation. Thus, we introduce the theoretical optimization framework, which the `R` package `SDT` is based on and that allows to compute the solutions to the afore mentioned issues. Section 4 presents the package `SDT` and we describe the functions of it. Section 5 demonstrates the package by examples and we apply its functions to an accompanying dataset named `learning_motivation`. In Section 6, we summarize and conclude with final remarks.

## 2. SELF-DETERMINATION THEORY

SDT provides a framework for the study of human motivation (Deci & Ryan, 1985, 2000, 2002; Ryan & Deci, 2000a). Empirical data corroborate that there are three basic psychological needs “essential in promoting life satisfaction and well-being”, the “opportunities to experience autonomy, competence, and relatedness” (Levesque, Zuehlke, Stanek, & Ryan, 2004, p. 68). Hereby, “autonomy refers to volition – the organismic desire to self-organize experience and behavior and to have activity be concordant with one’s integrated sense of self” (Deci & Ryan, 2000, p. 231). Competence refers to the universal want to control outcome and experience mastery (White, 1959), and “we consider competence or effectance to be one of the three fundamental psychological needs that can energize human activity” (Deci & Ryan, 2000, p. 231). “Relatedness refers to the desire to feel connected to others – to love and care, and to be loved and cared for” (Deci & Ryan, 2000, p. 231).

According to SDT, these three needs can be satisfied differently among individuals, but in any case their satisfaction is vital for the healthy development and well-being of all individuals (Deci & Ryan, 1985, 2000, 2002; Deci & Vansteenkiste, 2004). People can be moved by external factors (e.g., reward systems, grades, or evaluations) or be driven from within (e.g., interests, curiosity, or abiding values). Whereas the former examples are extrinsic, the latter examples are intrinsic and not necessarily externally rewarded or supported. SDT provides a basis for the study of the relationship between the extrinsic forces that externally act on humans and the intrinsic motives and needs internally inherent in humans.

Interpreted in a pedagogical context, the students’ motivational behavior will be more self-regulated if these basic psychological needs are better satisfied (Deci & Ryan, 1985, 2000, 2002; Ryan & Deci, 2000b, 2009; Müller, Hanfstingl, & Andreitz, 2007). The motivational behavior can be described by a continuous scale. According to Figure 1, this scale is segmented into intrinsic motivation, extrinsic motivation, and amotivation. Intrinsic motivation and amotivation are not further differentiated. Extrinsic motivation is separated into integrated regulation, identified regulation, introjected regulation, and external regulation. For details, see Deci and Ryan (2000, 2002).

Applications of SDT are numerous. An extensive reference list, including comprehensive materials on the theory and the questionnaires developed to assess the different SDT constructs, is available at http://www.selfdeterminationtheory.org.

Figure 1 displays the self-determination continuum. Thus, according to SDT, the behavior of a person can shift from extrinsic to intrinsically motivated. From left to right, the behavior is more and more internalized through the regulation types that are ordered along the continuum. Introjected regulation and identified regulation are relevant to the discussion of this paper. Introjected regulation refers to a person that is acting on the basis of external societal expectations only partially internalized and that remain external to the self. Identified regulation means the person has identified with the external values of his/her behavior and has internalized these more into her/his value system. Details can be found in Deci and Ryan (1985, 2000, 2002).

The subscales of external regulation and intrinsic regulation are, by theory, completely external motivation and internal motivation, respectively. For the intermediate subscales of introjected regulation and identified regulation, on the other hand, their internalizations are expressed as *“somewhat external”* and *“somewhat internal”*, respectively. That is, these intermediate regulation types are mixtures of external motivation and internal motivation and remain vaguely specified in SDT. The constrained regression analysis approach to quantifying these notions, along with the major implementation components, are theoretically presented in the following section.

## 3. CONVEX OPTIMIZATION AND MOTIVATION INTERNALIZATION

We start with a general introduction to convex optimization in the first paragraph of this section. But in the second and following paragraphs, it will be clear why we should care about convex optimization, meaning what problem convex optimization is going to solve in SDT.

Optimization is omnipresent in many different scientific fields and especially powerful if based on convexity (e.g., Boyd & Vandenberghe, 2009; Dattorro, 2009). The general problem of convex optimization can be stated as:

*minimize g*(π) *subject to* π ∈ 𝒟,
where *g*: ℝ^k^ → ℝ is a convex function mapping *k* arguments of interest
into a real-valued summary or target criterion,
and, determined by convex inequality (and affine equality) constraints,
𝒟 ⊂ ℝ^k^ is the convex set of all feasible values for the arguments.

The program is to minimize an objective function with respect to parameters of interest, under given side constraints on the parameters. The convexity assumptions for the objective function and the constraints ensure useful mathematical properties such as the characterization of (global) optimality based on the important in optimization Karush-Kuhn-Tucker conditions (Karush, 1939; Kuhn & Tucker, 1951; Kuhn, 1976; Roberts & Varberg, 1973; Hiriart-Urruty & Lemaréchal, 2001; Boyd & Vandenberghe, 2009).

An interesting and basic instance of this general convex optimization problem appears in SDT, related to the problem of motivation internalization. We apply optimization to gauge the internal and external motivation shares of the intermediate regulation types, presupposing that the relevant subscales have been measured using reliable and valid inventories. Each of the identified regulation and introjected regulation is modeled as a convex combination of the fully internal and fully external regulation types of intrinsic motivation and extrinsic motivation, respectively. The optimization problem implied is: to find for the identified and introjected regulation types those shares that minimize the discrepancy between the observed values based on the inventory scores and the values predicted by the convex combinations, having the unknown and to be estimated shares as their weights in the apparently extreme poles of the theory. There are two inequality constraints to consider, namely that the two shares in regards to intrinsic regulation and external regulation are nonnegative, and the equality constraint is that these shares must add up to 1.

For this outlined SDT convex optimization problem, which is a basic one, the question can be phrased as a quadratic program. This means that we can have a special convex quadratic form for the objective function, with corresponding affine inequality and equality constraints, which then makes possible the application of readily available numerical algorithms for efficiently solving the program. For this purpose, we will use the method by Goldfarb and Idnani (1982, 1983). The latter is a numerically stable dual method for computing the solutions of quadratic programming problems of the type we encounter in this paper.

Let *X*_1_ = *InR* and *X*_2_ = *ExR* be the intrinsic regulation (*InR*) and external regulation (*ExR*) types, which are assumed to be internal and external, respectively. Let *Y* stand for either identified, *IdR*, or introjected, *IjR*, regulation, for which we want to compute the internalization or externalization shares. The basic model is

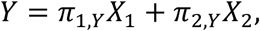

where *π*_1,Y_ ≥ 0, *π*_2,Y_ ≥ 0, and + *π*_1,Y_ = 1, and *Y*, *X*_1_, and *X*_2_ stand for the data and are taken over all sample units (e.g., students). The parameters *π*_1,Y_ and *π*_2,Y_ are unknown and estimated from the data. In other words, *IdR* and *IjR* can be modeled as a convex combination of *InR* and *ExR*. That is, the degree of internalization is gauged by the shares *π*_1,*Y*_, as the internal extent of identified or introjected regulation, and *π*_2,*Y*_, as the external extent of identified or introjected regulation, relative to the extreme internal and external poles of the self-determination continuum (cf. Figure 1).

The extension of this model to more than two components is straightforward:

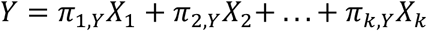, where *π*_*i,Y*_ ≥ 0 for 1 ≤ *i* ≤ *k*, and ∑*π*_*i,Y*_ = 1.

Subsequently, we consider the general formulation and omit the subscript *Y*, having in mind that one of both *IdR* or *IjR* (or any other SDT target variable) is being considered. The *X*_*i*_’s for 1 ≤ *i* ≤ *k* form what we call the reference system of base elements, according to which the convex decomposition of the target variable *Y* is made, with the *π*_*i*_’s for 1 ≤ *i* ≤ *k* interpreted as the corresponding shares in this system.

A special choice of the target variable and reference system can be made for the analysis of the motivation simplex structure posited by SDT. (For example, the target variable can be intrinsic regulation, and the reference system can consist of identified regulation, introjected regulation, and external regulation. This choice is exemplified in Section 5). Ünlü and Dettweiler (2015, p. 685): “The *simplex structure* of self-determination theory means that motivation regulation types theoretically closer to one another are more strongly interrelated, indicating that the self-determination theory regulatory styles can be linearly ordered along the underlying continuum (Ryan & Connell, 1989; Deci & Ryan, 2000).” In the SDT literature, “interrelated” is synonymous with “correlated”, and Ünlü and Dettweiler (2015) have proposed assessing that structure based on optimal shares instead. Thus, under a simplex structure assumption, we expect in this new approach that the computed shares are larger for motivation regulation types theoretically closer to one another.

A numerical solution to the optimization problem raised in SDT can be derived as follows. Formulated in analogy to the general convex optimization problem, we

*minimize*

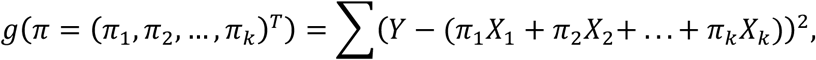

where “.^*T*^” stands for the transpose of a matrix and the sum is taken over all sample units (e.g., students),

*subject to*

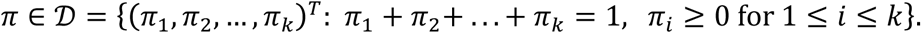

There are *k* inequality constraints and one equality constraint. It can be proven that the target function *g* is convex and that the feasible set 𝒟 is convex (and even compact).

This problem can be viewed as a quadratic program. Obviously, an equivalent formulation of the problem is:

*minimize*

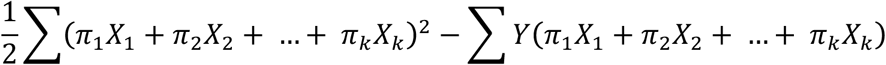

*subject to*

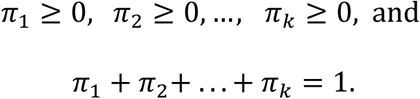

This can be written in matrix notation yielding the required quadratic program expression. More precisely, the first term is equal to

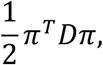

where *D* = (*X*_1_, *X*_2_,…, *X*_k_)^*T*^ (*X*_1_, *X*_2_,…, *X*_k_) and the SDT variables *X*_1_, *X*_2_,…, *X*_k_, and *Y* are column vectors and observed data. The second term above equals

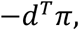

where *d* = (*X*_1_, *X*_2_,…, *X*_k_)^*T*^ *Y* and the surveyed SDT variables are used as column vectors in this notation. Moreover, the *k* inequality constraints can be written as

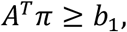

where *A* = *I_k_* is the *k*×*k* identity matrix, and *b*_1_ = 0_*k*_ is the column vector of length *k* containing only 0’s. The equality constraint is *a*^*T*^ *π* = *b*_2_, where *a* = 1_*k*_ is the column vector of length *k* consisting of 1’s only, and the scalar *b*_2_ = 1.

In sum, this yields the required (convex) quadratic program that corresponds to our initial SDT question:

*minimize*

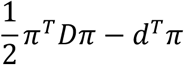

*subject to*

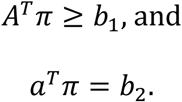

Given the quadratic form above, we can use software to calculate a solution. This is implemented in the `R` package SDT, described in Section 4.

The computable shares not only can be used for internalization or simplex structure analyses, but can also provide an adjusted self-determination or relative autonomy index, SDI or RAI. Scoring protocols such as the original SDI or RAI index are summary statistics that aggregate test scores to give an overall informative measure (see Grolnick & Ryan, 1989; Ryan & Connell, 1989; Vallerand, 2007; Wilson, Sabiston, Mack, & Blanchard, 2012; Chemolli & Gagné, 2014).

The original SDI or RAI index is defined as

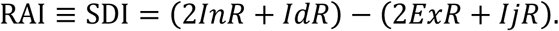

This scoring protocol does not take into consideration the fact that the identified and introjected regulation types are mixtures of internal and external motivation. The resulting overall measure may be confounded and therefore may lack interpretability, because in weighting the subscale scores the same weights are used for the two shares of internal and external motivation of a regulation type.

The adjusted SDI or RAI (see Ünlü, 2016), which is weighted according to the extent to which these regulation types are internal and external, is given by

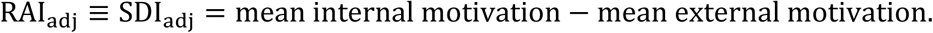

Mean internal motivation, 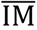, and mean external motivation, 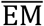, are quantified using the *π*-weights obtained from the quadratic program described above (with identified regulation or introjected regulation as the target variable, and with intrinsic regulation and external regulation as the reference system):

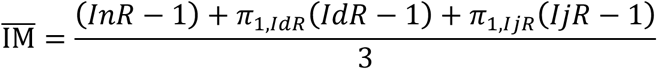

and

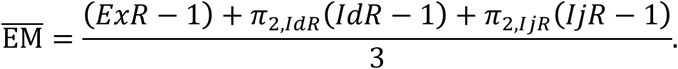

Translation with −1 and averaging guarantee that all of the instrument variables *InR* − 1, *IdR* − 1, *IjR* − 1, *ExR* − 1, the components 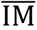 and 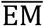, and the new scoring protocol RAI_adj_ ≡ SDI_adj_ range in the same interval. This is not true for the original index. In the `R` package `SDT`, both indices are implemented, described in the following section.

## 4. R PACKAGE SDT

We briefly describe the functions and relevant parts of the package. How to actually use the software is demonstrated by examples in Section 5. The description of the package will be short, and detailed information can be found in the comprehensive documentation files and commented source code for the package in `R` (for an overview, type `package? SDT`).

The package `SDT` uses the `S3` system and consists of the following main functions: `internalization` (motivation internalization analysis), `sdi` (original and adjusted SDI or RAI index), and `simplex` (motivation simplex structure analysis). It contains further functions, which are `plot`, `print`, and summary methods: `plot.sdi`, `print.sdi`, and `summary.sdi`, and `plot.share` and `print.share`. There is the accompanying dataset `learning_motivation` (described and analyzed in Section 5), based on which the features of the package `SDT` are illustrated.

One of the main functions of the package is `internalization: internalization(intermediate_regulation, intrinsic_regulation, external_regulation)`

This function provides the motivation internalization or externalization computation (see Section 3). It takes an intermediate regulation type, either identified or introjected, as the target variable, and returns its shares, a numeric vector containing two named components `internal` `share` and `external share`, with respect to the poles of intrinsic regulation and external regulation as the reference system. The arguments `intermediate_regulation`, `intrinsic_regulation`, `external_regulation` are numeric vectors of respective subscale motivation scores, where no infinite, undefined, or missing values are allowed.

The original and adjusted SDI or RAI indices can be computed using the function

~~~
sdi:
sdi(intrinsic_regulation, identified_regulation, introjected_regulation, external_regulation, compute.adjusted = TRUE, minscore = 1)
~~~

This function takes as input the four regulation types, which are numeric vectors of intrinsic, identified, introjected, and external regulation subscale motivation scores, respectively, where no infinite, undefined, or missing values are allowed. The argument `compute.adjusted` = `TRUE` indicates adjusted index computation, whereas specifying `FALSE` for it allows to compute the original index. The argument `minscore` gives the minimum score of the scale procedure (typically 1) and only needs to be specified for the adjusted index (for `compute.adjusted = FALSE`, this argument is irrelevant and ignored). As mentioned in Section 3, for the adjusted variant, we translate with “− `minscore`” and average to warrant that all variables and component and index values range in the same interval.

The function `sdi` returns a named list. The returned list contains three components, in both the cases of original or adjusted index computation. For the original index computation, the components `confounded_internal_locus, confounded_external_locus`, and `sdi_original` are numeric vectors of the confounded internal locus values (*2InR* + *IdR*), confounded external locus values (*2ExR* + *IjR*), and the overall original index values, for all students or rows of the dataset. For the adjusted index computation, the components `adjusted_internal_locus, adjusted_external_locus, and sdi_adjusted` are numeric vectors of the adjusted internal locus values 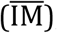, adjusted external locus values 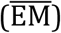, and the overall adjusted index values, for all students or rows of the dataset.

For simplex structure analysis, the function `simplex` can be used:

~~~
simplex(target_regulation, base_regulation_1, base_regulation_2, base_regulation_3)
~~~

The simplex structure shares are calculated of a target regulation type, either intrinsic, identified, introjected, or external regulation subscale motivation scores, in the reference system of the remaining base regulation types. In these numeric vectors, no infinite, undefined, or missing values are allowed. The function `simplex` returns a numeric vector consisting of three components: `base_regulation_1 share, base_regulation_2 share, and base_regulation_3 share`; these are the respective shares of the target regulation relative to the remaining base regulation types of the theory.

The interdependencies among the functions are as follows. In the functions `internalization` and `simplex`, the function `solve.QP` of the `R` package `quadprog` (`S` original by Berwin A. Turlach `R` port by Andreas Weingessel, 2013) is used, to solve the SDT related convex quadratic program (see Section 3). `solve.QP` implements the quadratic program minimizer by Goldfarb and Idnani (1982, 1983). For calculation of the adjusted index, the function `sdi` calls the function `internalization`.

The functions `internalization` and `simplex` return objects (denoted x) of the class “`share`”. For these, `S3` `plot` and `print` methods are implemented. The `plot` method

`plot(x, target = NULL, reference = NULL,…)`

graphs the results of SDT share analyses by means of stacked bar plots of the internalization, externalization, or simplex structure shares of a target regulation relative to a reference system. Generic or user-specified labeling of the plot axes are possible. The default values `target = NULL` and `reference = NULL` correspond to generic labeling. If in user-specified labeling character strings for the arguments `target or reference` are specified, these are used to label the *x*-axis and *y*-axis of the bar plot, respectively. What this means is also shown by examples in Section 5, and detailed information on the labeling can also be found in the comprehensive documentation files in `R`. The `print` method

`print(x,…)`

outputs on the console the shares of internalization, externalization, or simplex structure, stripped off the attributes.

The function `sdi` returns an object (`x or object`) of the class “`sdi`”. There are corresponding `S3 plot, print`, and summary methods. The `plot` method

`plot(x, minscore = 1, maxscore = 5,…)`

visualizes the results of the original or adjusted SDI or RAI index computation. A scatterplot is drawn of the confounded or adjusted external locus values on the *y*-axis, and the confounded or adjusted internal locus values on the *x*-axis. The line *y = x* is shown as the red full line, for graphical comparison of the two value types. The admissible range for the original or adjusted component values is displayed in gray dashed lines. Points with larger overall index values are portrayed in darker gray tone. The `minscore` and `maxscore` arguments are used to define the admissible range. `minscore` is the minimum score of the scale procedure (typically 1). `maxscore` is the scale procedure maximum score (typically 4, 5, or 7). For the adjusted index, the admissible range is [0, `maxscore` − `minscore`], with [0, 4] in the default values. The admissible range for the original index is

[(2 ∙ `minscore`) + `minscore`, (2 ∙ `maxscore`) + `maxscore`], which is [3, 15] for the default values.

The print method

`print(x,…)`

prints the original or adjusted overall SDI scores, for all students or rows of the dataset.

The summary method

`summary(object,…)`

outputs simple summary statistics for the confounded or adjusted internal locus component values, confounded or adjusted external locus component values, and for the values of the original or adjusted SDI overall index. The summary statistics printed are the minimum, first quartile, median, mean, third quartile, and the maximum.

## 5. EXAMPLES

The goal of this section is to illustrate by examples how the functions of the package can be run technically. As such, following the here described use cases, the functions can be analogously applied in any other empirical data set. This section simply shows how. In particular, the goal of this paper cannot be to systematically investigate and research, centered around real-world applications, the scope and limitations of the techniques of SDI adjustment, simplex structure analysis, and motivation internalization, from a substantial point of view. This is more out of the scope rather than a limitation of this software paper. The present paper provides the software basis for such substantial future research work.

### 5.1. Dataset

The package SDT contains a real dataset, on learning motivation from Austrian school classes in mathematics, information sciences, and natural sciences (Müller et al., 2007): `learning_motivation`. (I would like to thank Professor Dr. Florian Müller and his colleagues from University of Klagenfurt, Austria for providing the author with this dataset.) We use this dataset to illustrate the package’s functions. The `learning_motivation` data frame consists of 1,150 rows/students and 6 columns/variables. The students comprise 578 girls and 572 boys (mean age 14.1, with standard deviation 1.9). The variables are sex (integer vector, female = 1, and male = *2*), age (integer vector, years), and the learning motivation scores for the subscales of intrinsic regulation, identified regulation, introjected regulation, and external regulation. The motivation variables of the data frame are numeric vectors, which contain aggregate subscale scores, that is, the means taken over all test items that form a respective subscale.

The first six rows of the data frame are (R input is marked as “R>”, and the corresponding `R` output is shown below the input):

`R> head(learning_motivation)`

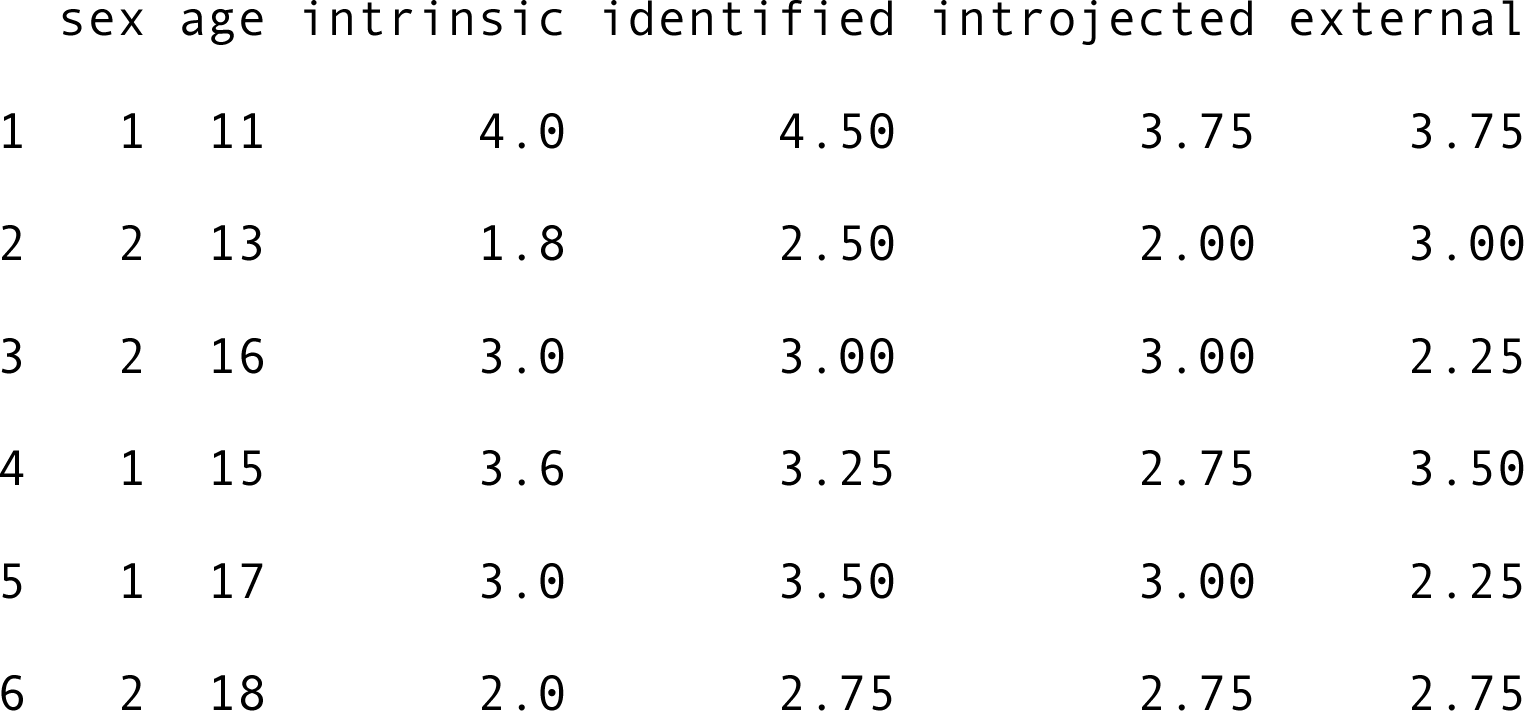

We attach the data frame to the search path of the `R` session

`R> attach(learning_motivation)`

so in subsequent analyses we can easily access any variable of the dataset directly by typing its name.

### 5.2. Internalization Analysis

We have the intermediate motivation variables identified and introjected. To compute their shares of internalization or externalization, we run the function internalization on the variables:

`R> (idr <- internalization(identified, intrinsic, external)) internal share external share`

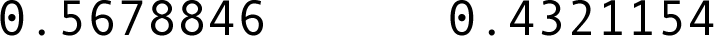

`R> (ijr <- internalization(introjected, intrinsic, external)) internal share external share`

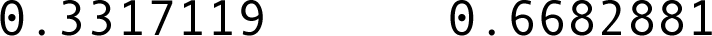

Identified regulation is composed of approximately 57% internal share and 43% external share, which is more internal motivation than external motivation, as expected by theory. For introjected regulation, which according to theory ought to be more external motivation than internal motivation, we have the internal and external shares of approximately 33% and *67%*, respectively.

We can access the attribute value and class of the object idr, or print all attributes of the object ijr:

`R> attr(idr, "analysis")`

`[1] "internalization"`

`R> class(idr)`

`[1] "share"`

`R> attributes(ijr)`

`$names`

`[1] "internal share" "external share"`

`$analysis`

`[1] "internalization"`

`$class`

`[1] "share"`

Objects such as ijr of the class “`share`” can be plotted:

`plot(ijr)`

gives the stacked bar plot with generic labels of the axes shown in Figure 2.

**Figure 2.**
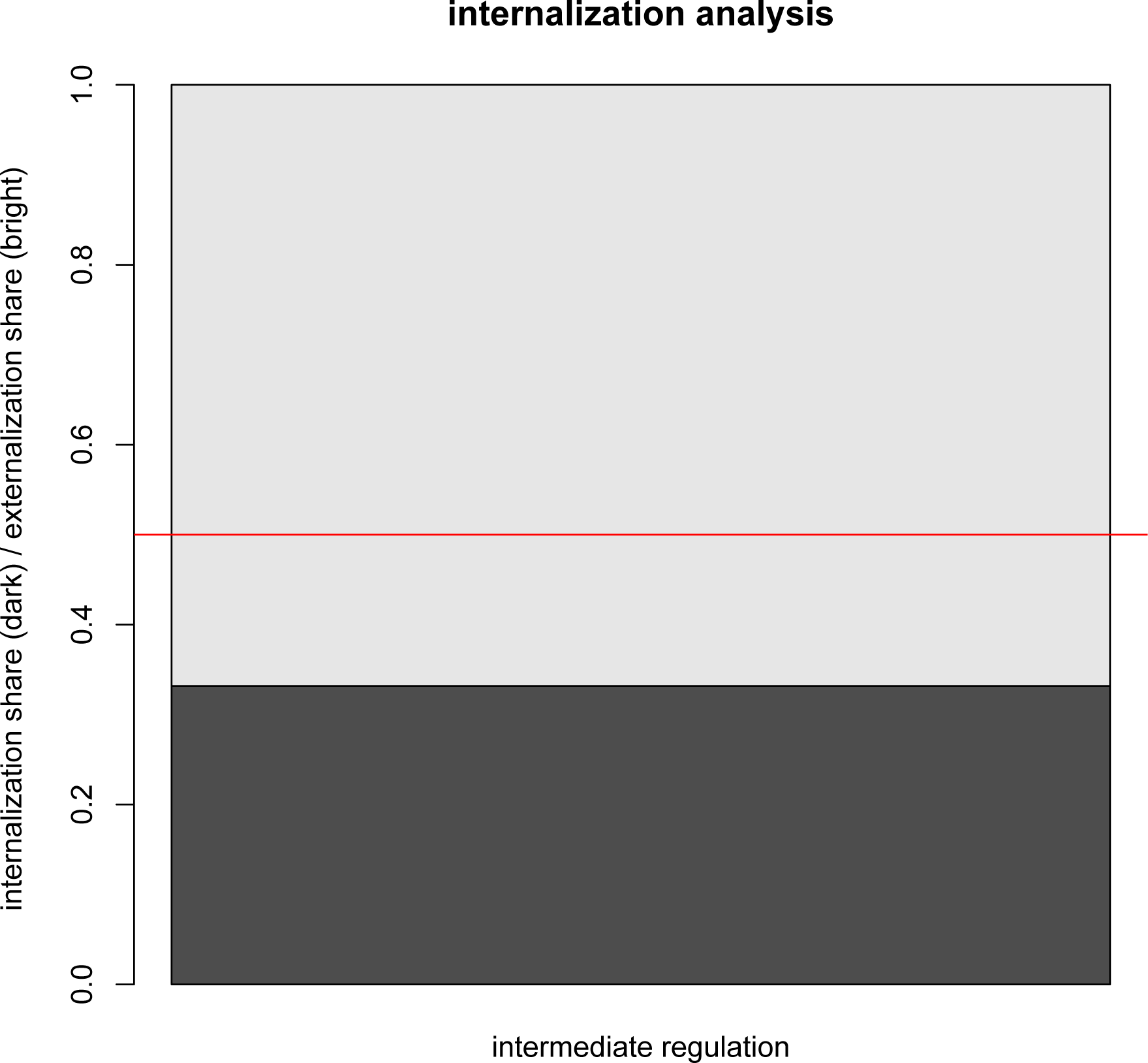
Internalization analysis plot for introjected regulation as the target variable. Stacked bar plot is shown with generic labels.

We can have a similar plot for the object idr with user-specified labels

`plot(idr, target = "identified regulation", reference = c("intrinsic regulation", "external regulation"))`

which is shown in Figure 3.

**Figure 3.**
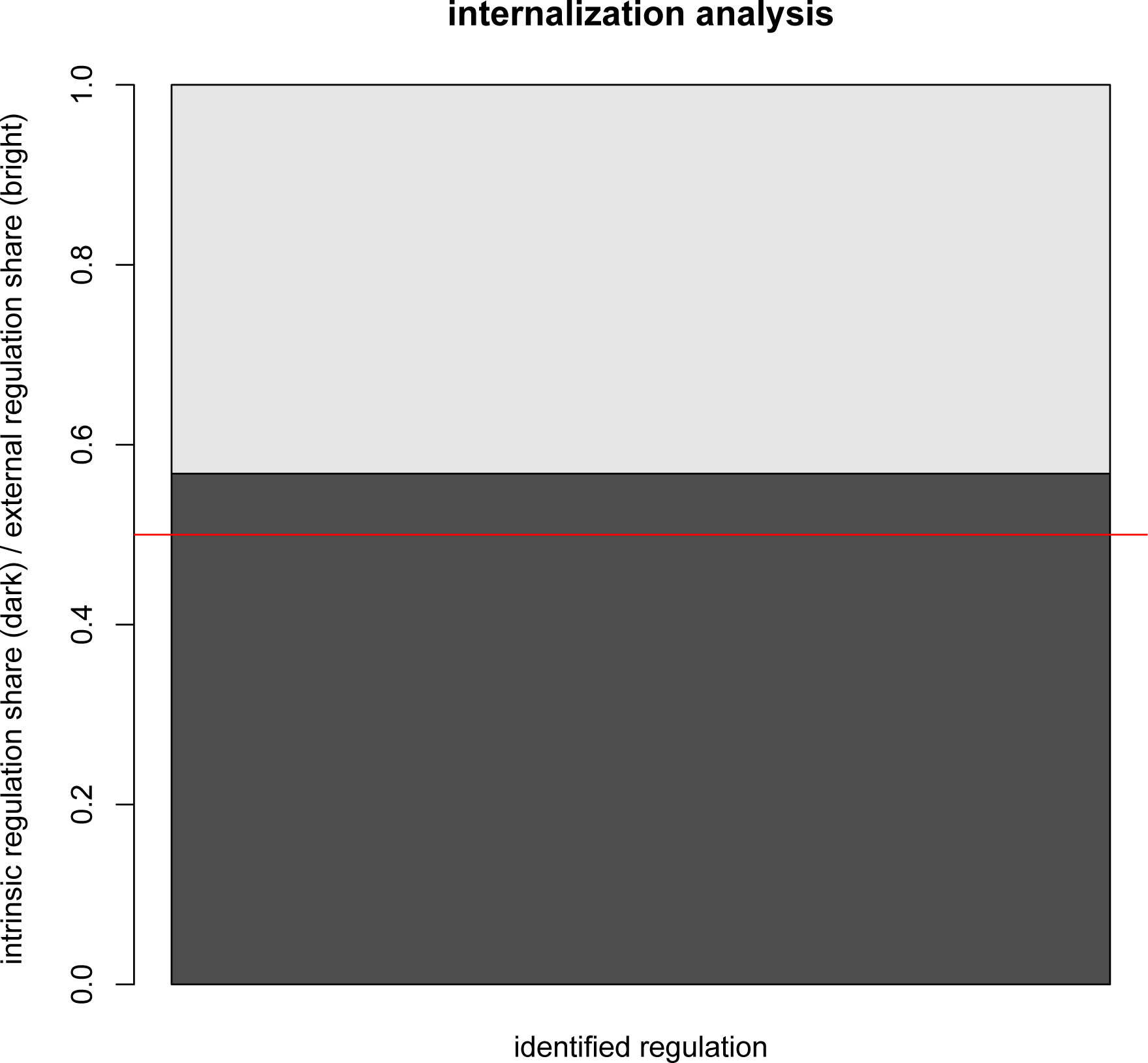
Stacked bar plot of the internal and external shares of identified regulation. User-specified labels are provided.

### 5.3 Simplex Structure Analysis

We can perform a simplex structure analysis with intrinsic regulation as the target variable, and with identified regulation, introjected regulation, and external regulation as the reference system:

`R> (simstr <- simplex(intrinsic, identified, introjected, external)) base_regulation_1 share base_regulation_2 share base_regulation_3 share`

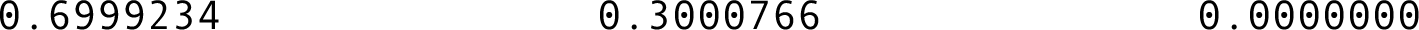

We can see that the posited simplex structure assumption is fulfilled for this choice of variables. The computed shares are plausible with theory. Intrinsic regulation, which is completely internal, is more interrelated with identified regulation with a share of approximately 70%, followed by introjected regulation with a share of approximately 30%, and has a 0% share in regard to external regulation, which is completely external.

The object simstr is a numeric vector with an attribute value and class:

`R> mode(simstr)`

`[1] "numeric"`

`R> attr(simstr, "analysis")`

`[1] "simplex"`

`R> class(simstr)`

`[1] "share"`

and can be plotted with user-specified labels

`R> plot(simstr, target = "intrinsic regulation", reference = c("identified regulation", "introjected regulation", "external regulation"))`

shown in Figure 4, where the external regulation share of the computed value zero is omitted.

**Figure 4.**
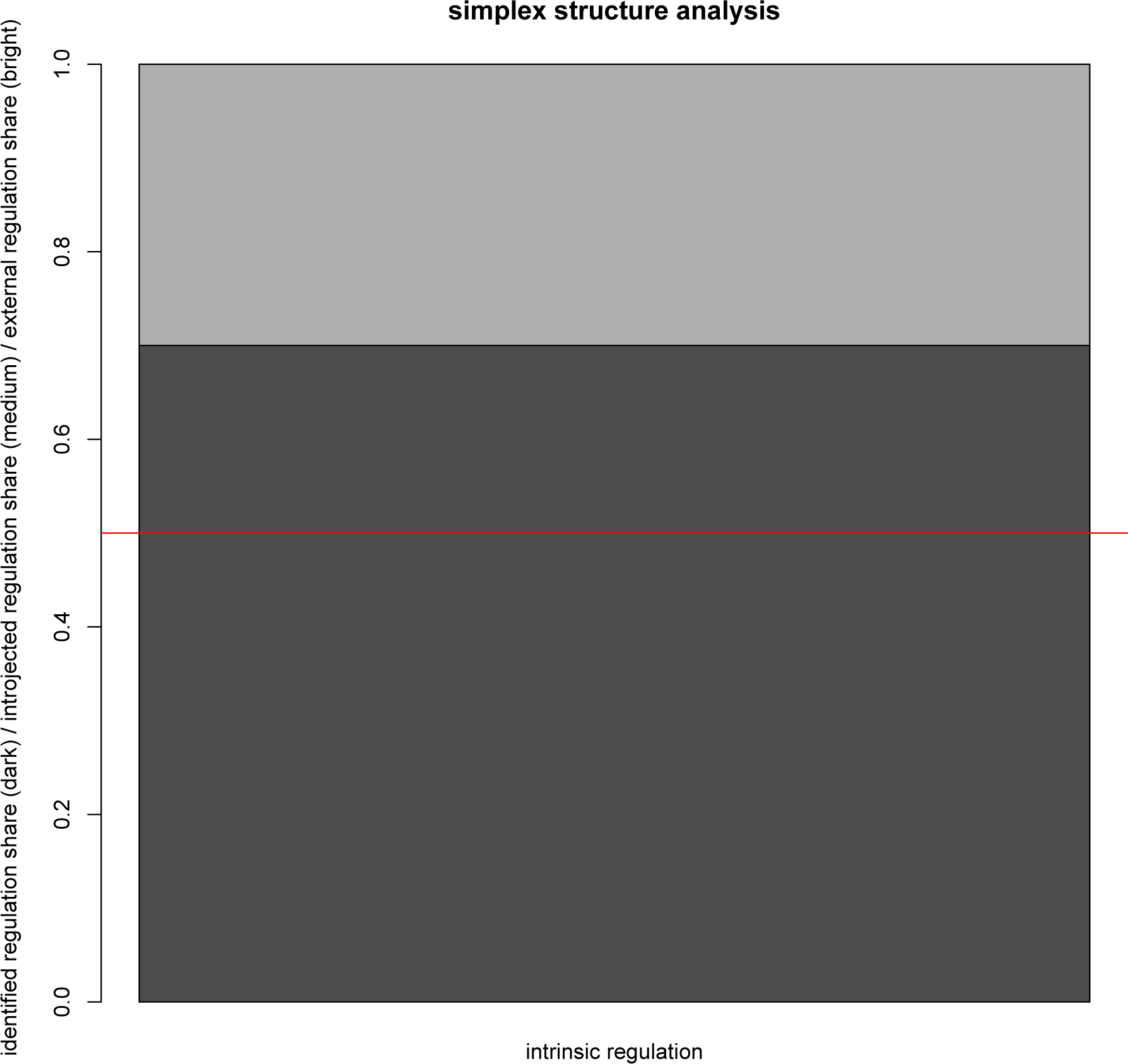
Simplex structure analysis plot, with user-specified labels, for intrinsic regulation as the target variable, and with identified regulation, introjected regulation, and external regulation as the reference system.

A similar plot can be produced with external regulation as the target variable, and with intrinsic regulation, identified regulation, and introjected regulation as the reference system, with generic labels:

`R> plot(simplex(target_regulation = external,`

`base_regulation_1 = intrinsic,`

`base_regulation_2 = identified,`

`base_regulation_3 = introjected))`

This is shown in Figure 5, where the intrinsic regulation share of value zero is omitted.

**Figure 5.**
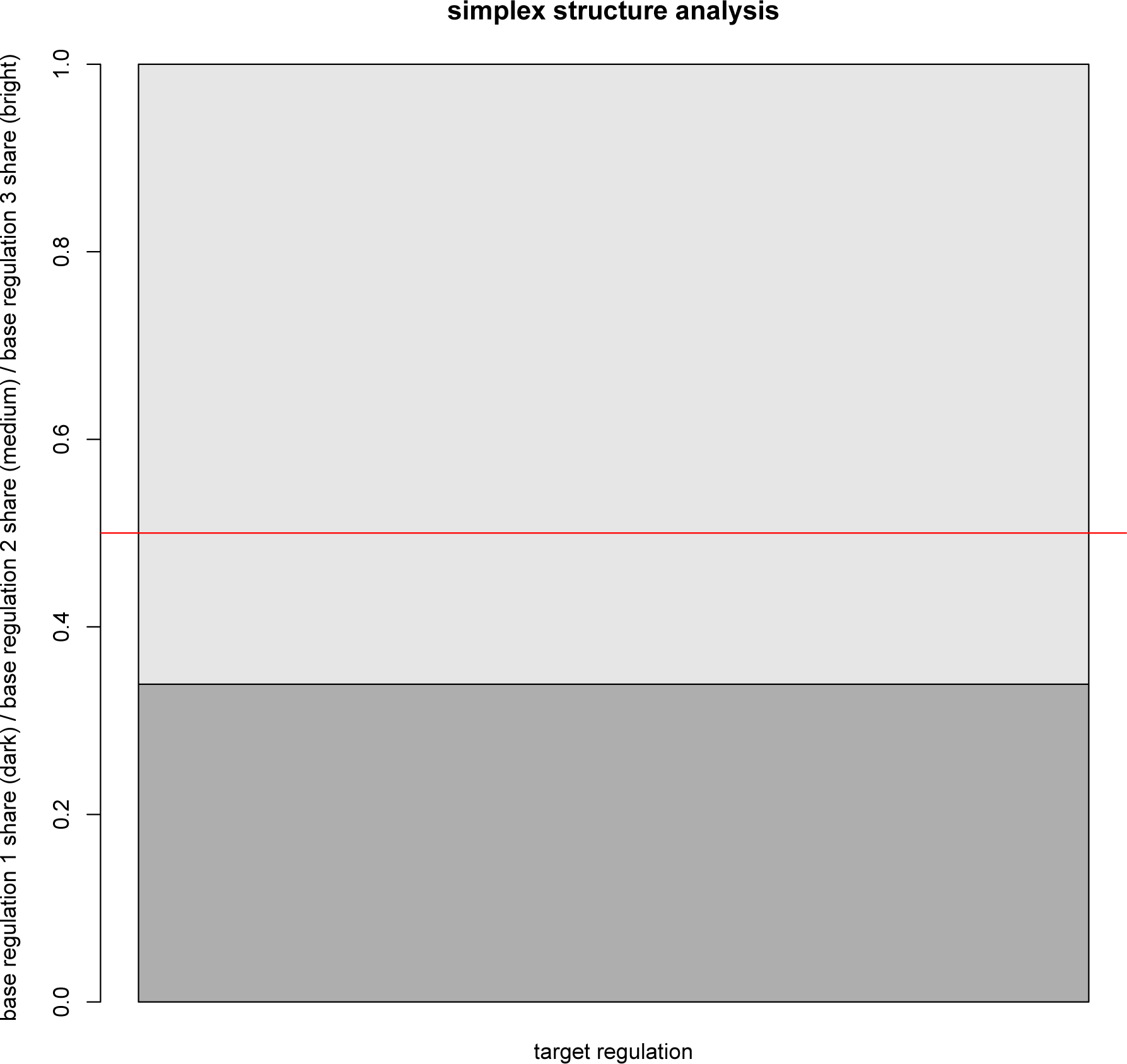
Stacked bar plot of the simplex structure shares of external regulation with respect to intrinsic regulation, identified regulation, and introjected regulation, displayed with generic labels.

The respective shares in this case are:

`R> simplex(external, intrinsic, identified, introjected) base_regulation_1 share base_regulation_2 share base_regulation_3 share`

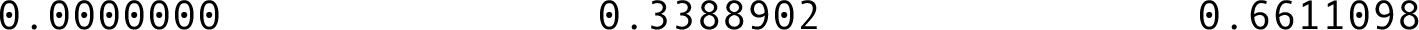

Again, the shares are in accordance with theory, and the simplex structure assumption is satisfied with this set of variables. External regulation is most interrelated with introjected regulation (66%), followed by identified regulation (34%), and with a zero share regarding intrinsic regulation.

### 5.4. Original and Adjusted Indices

We can compute, for each student or row pattern of the dataset, the corresponding values of the original and adjusted SDI or RAI indices, using the function sdi. Adjusted index computation can be performed by

`adj <- sdi(intrinsic, identified, introjected, external)`

with the attributes

`R> attributes(adj)`

`$names`

`[1] "adjusted_internal_locus" "adjusted_external_locus" "sdi_adjusted"`

`$variant`

`[1] "adjusted"`

`$class`

`[1] "sdi"`

We can inspect the first six elements of each list component vector.:

`R> lapply(adj, head)`

`$adjusted_internal_locus`

`[1] 1.9666ʘ13 ʘ.6611796 1.2663977 1.486ʘ787 1.361ʘ451 ʘ.8580979`

`$adjusted_external_locus`

`[1] 2.ʘ33399 1.1ʘ5487 1.15ʘ269 1.547255 1.222288 1.225235`

`$sdi_adjusted`

`[1] -ʘ.ʘ6679746 -ʘ.4443ʘ747 ʘ.11612865 -ʘ.ʘ6117589 ʘ.13875686`

`-ʘ.36713743`

The original index computation can be performed by

`orig <- sdi(intrinsic, identified, introjected, external, compute.adjusted = FALSE)`

and summarized using the corresponding method

`R> summary(orig)`

summary of confounded internal locus values: Min. 1st Qu. Median Mean 3rd Qu. Max.

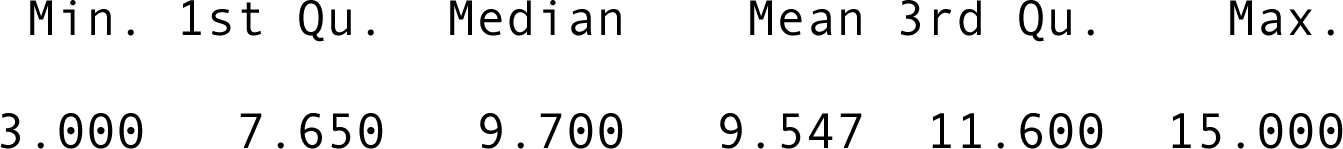

`summary of confounded external locus values:`

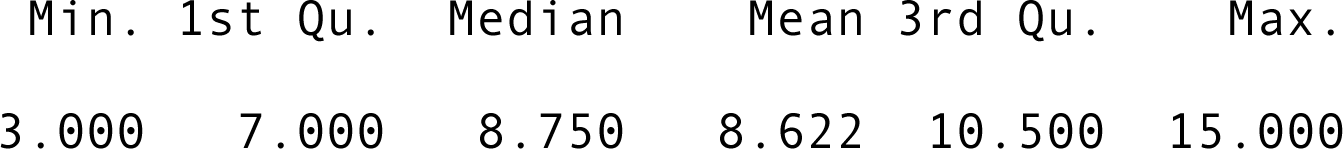

summary of original SDI scores:

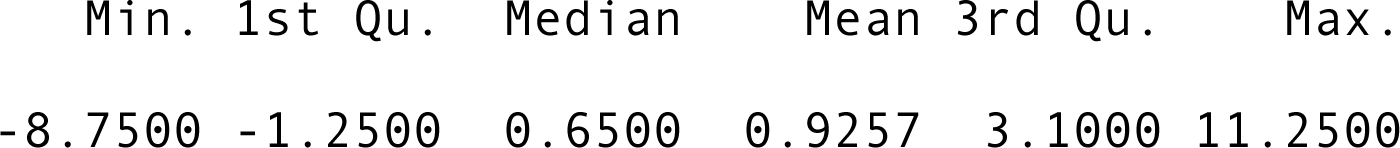

In contrast, the summary for the adjusted measure yields

`R> summary(adj)`

`summary of adjusted internal locus values:`

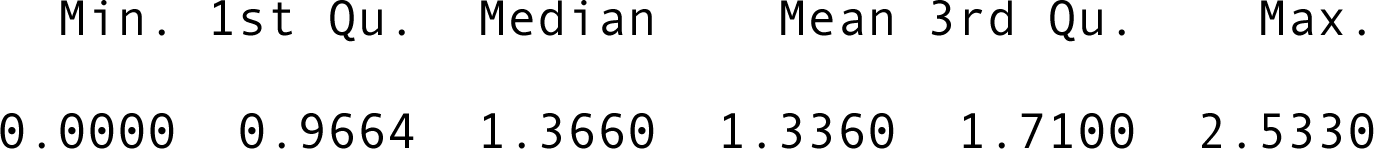

`summary of adjusted external locus values:`

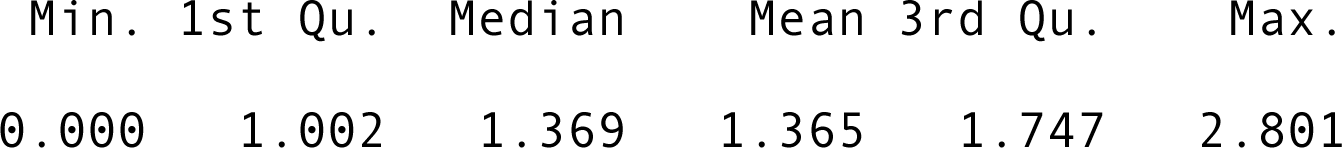

`summary of adjusted SDI scores:`

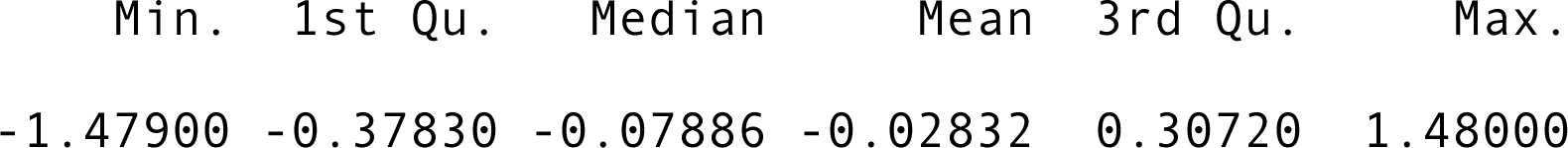

It is interesting to plot objects of the class “sdi”. Plotting the objects adj and orig (minimum and maximum scores of the scale procedure were the default values 1 and 5, respectively)

`plot(adj)`

`plot(orig)`

produce the scatterplots shown in Figures 6 and 7, respectively.

**Figure 6.**
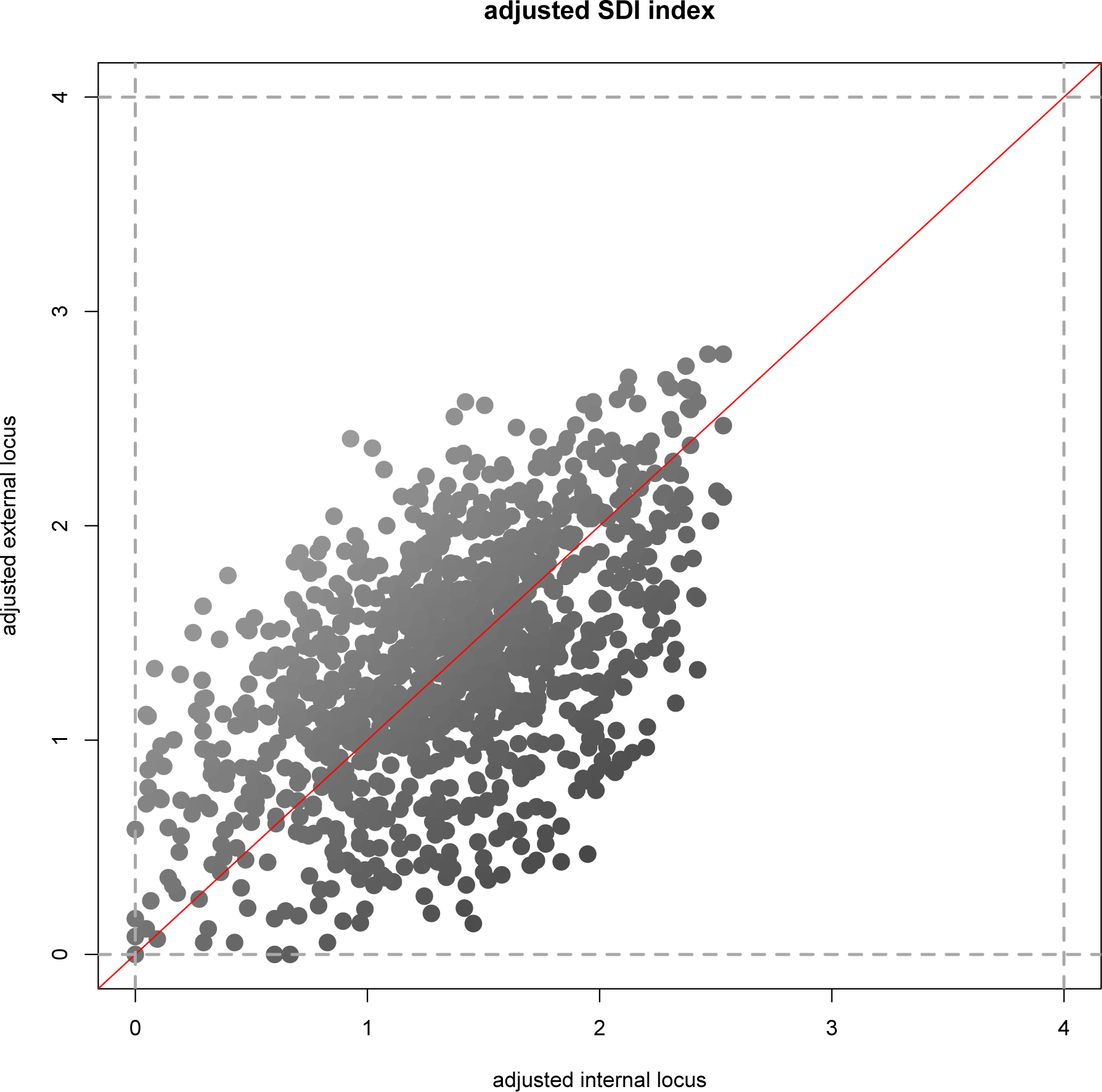
Scatterplot for the adjusted self-determination index (SDI). Adjusted external locus values versus adjusted internal locus values are plotted. The points in the scatterplot are shown at different gray levels, determined by their adjusted SDI overall index values. The red line, to assist visualization, is *y = x*, and the admissible range [0, 4] is graphed in gray dashed lines.

**Figure 7.**
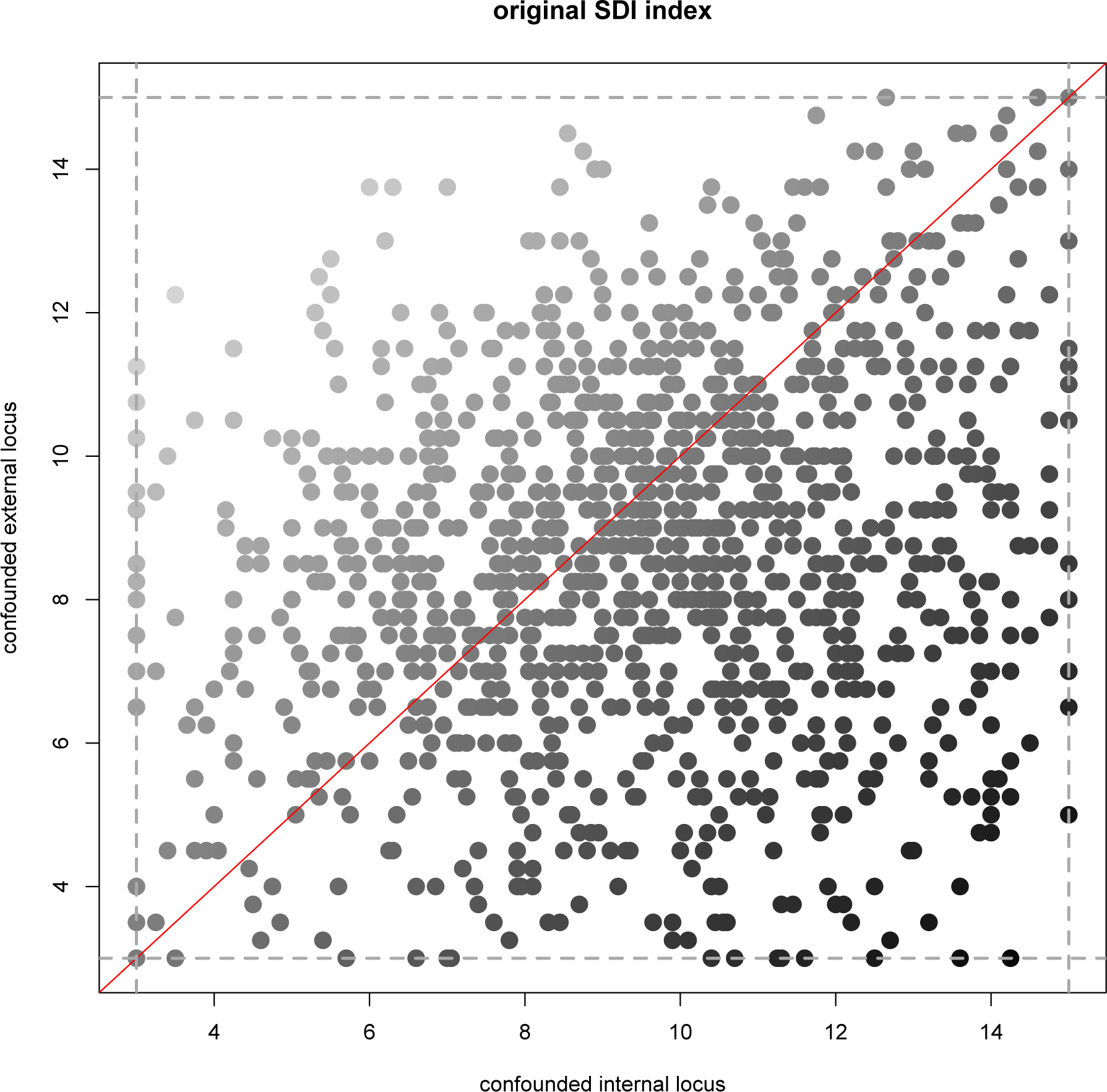
Scatterplot for the original self-determination index (SDI). Confounded external locus values versus confounded internal locus values are plotted. Points are drawn at different gray levels, depending on the original SDI overall index values. The red line, for comparison, is *y = x*, and the admissible range [3, 15] is shown in gray dashed lines.

Points with larger overall index values are depicted in darker gray tone. The admissible ranges for the original and adjusted indices are [3, 15] and [0, 4], respectively.

From Figures 6 and 7, we can see that the adjusted scores are more concentrated around the diagonal line shown in red, whereas the confounded scores do scatter messily over the broad range of admissible values. The adjusted scores in Figure 6 indicate that the external and internal motivation extents are distributed primarily in the lower to middle regions, between 1 to 2 scale points. This renders possible to see, if and where on the common scale from 0–4, there may be tolerance or scope for possible interventions, to improve on pupils’ learning motivation. This is not possible for the original index, which messily scatters.

Moreover, according to the adjusted RAI index, the girls are slightly extrinsically oriented:

`R> mean(adj$sdi_adjusted[sex == 1])`

`[1] -ʘ.1062873`

In contrast, with regard to the confounded original index, the girls can be deemed clearly intrinsically motivated:

`R> mean(orig$sdi_original[sex == 1])`

`[1] ʘ.3730969`

The former observation based on the adjusted index is more plausible. For, mathematics, informatics, and natural sciences school classes are studied, and there is empirical evidence that girls in these subject areas may typically behave extrinsically motivated.

The print method lists the original and adjusted SDI overall index values, for all students or rows of the dataset (the `R` output is omitted, for typographic reasons):

`R> adj`

adjusted SDI scores:

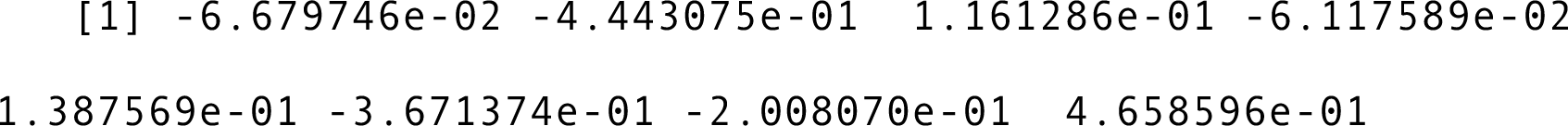

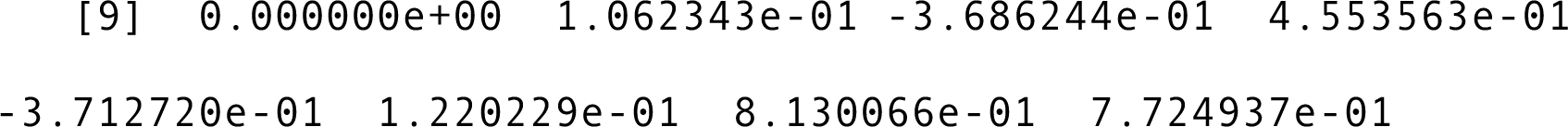

R> orig

`original SDI scores`:

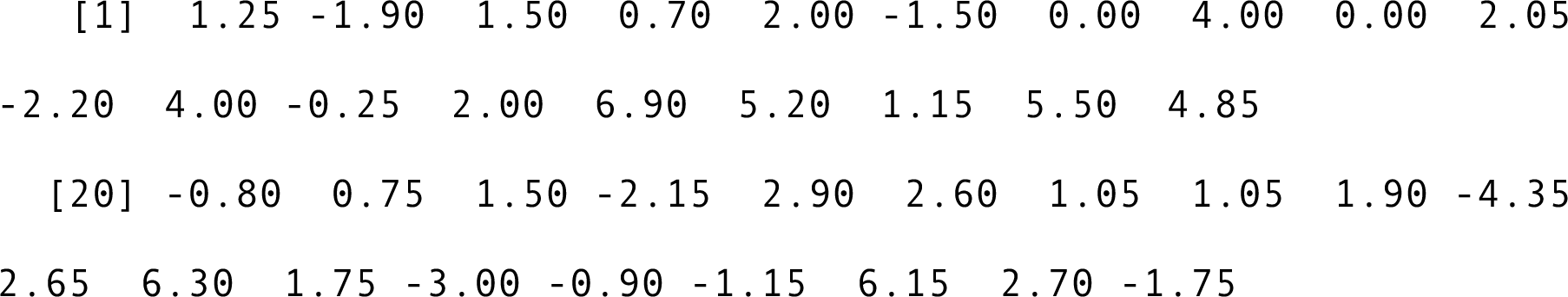

## 6. CONCLUSION

We have introduced the package SDT for computing self-determination theory (SDT) measures in the `R` language and environment. The package contains functions for computing the measures of motivation internalization, motivation simplex structure, and the original and adjusted self-determination or relative autonomy indices (SDI or RAI). The functions of the package SDT were described, and we demonstrated the functions’ usage on an accompanying example dataset.

With the package SDT in `R` we hope to have established a basis for computational work in SDT. We plan to extend this package to incorporate such dimensionality reduction approaches as principal component analysis and factor analysis, for SDT questionnaire validation in R. Interactive visualization techniques for the exploration of raw-data motivation variables could also be implemented and utilized in R, for exploratory SDT analyses. The realization of SDT, for the first time in R, can also be valuable in applying current or computational statistical methods to SDT. For instance, the determination of confidence intervals and hypothesis testing in SDT for the computed optimal shares and the original and adjusted SDI or RAI indices may likely be realized using resampling methods. Future work of this sort would involve extensive computer simulation, which could be ideally achieved with R.

